# The lack of simplicity in sequence-fitness relationships

**DOI:** 10.64898/2026.04.30.722031

**Authors:** Kristina Crona, Devin Greene

## Abstract

Gene interactions play an important role in the development of antimicrobial drug resistance and other evolutionary processes of medical importance. Empirical studies have revealed multiple peaks, inaccessible trajectories, and constraints on mutation order. Higher order epistasis is associated with obstacles in fitness landscapes. However, its significance has been debated in recent years, sometimes through reinterpretations of data from previous publications.

We suggest that local higher order interactions may help reconcile these seemingly contradictory findings. Rank order based methods can be informative when other methods fail to detect consequential interactions. In addition to conventional rank order methods, including sign epistasis, we introduce signed bipyramids. A bipyramid interaction compares extreme genotypes against their intermediates, for example a triple mutant and the wild-type against the corresponding single mutants. In general, interactions are signed if they are implied by the rank order alone.

## 1. Introduction

The role of higher order epistasis in evolutionary biology has long been debated (Weinrech et al., 2013). One can find older theories that claim that the fitness of a genotype is determined by main effects and pairwise epistasis only. If correct, then the fitness of arbitrary genotypes would be determined by the wild-type, single mutants, and double mutants only. Such theory can safely be rejected based on years of modern sequencing data.

A less radical proposal is that many phenotypic measures can, to a high degree of accuracy, be described without incorporating higher order effects, provided one allows for global epistasis (Park et al., 2024). Specifically, the proposed approach applies a sigmoid link function for describing global nonlinearities. Other approaches to global non-linearities include Otwinski et al. (2018); Starr et al. (2016); Kryazhimskiy et al. (2014).

However, empirical landscapes reveal features that seem at odds with the claimed simplicity. For example, some antimicrobial resistance landscapes have multiple suboptimal peaks and evolutionarily inaccessible trajectories to the peaks. The constraints can force an adapting population to accumulate mutations in a specific order, thereby slowing the emergence of resistance. In some cases, these features create opportunities for treatment strategies that exploit the intrinsic barriers of the landscape (Nichol et al., 2019; Mira et al., 2015).

There is an association between many peaks and higher order epistasis, based on an exhaustive search of 93,270,310 4-cube graphs (Crona et al., 2020), and similarly for 3-cube graphs. In particular, the graph in Figure 1, where each arrow points toward the genotype of higher fitness, is incompatible with the absence of higher order interactions. The number of peaks, i.e., sinks in the graph, exceeds what is possible for a biallelic 3-locus system with no higher order interactions. Figure 2 shows malaria drug resistance and Figure 3 antibiotic resistance. Here we show that these graphs are incompatible with the absence of higher order interactions by applying theory on partial orders and higher order interactions (Crona et al., 2020; Lienkaemper, 2018; Crona et al., 2017). Informally, there is a limit to how hostile a fitness landscape can be in the absence of higher order epistasis.

**Figure 1.**
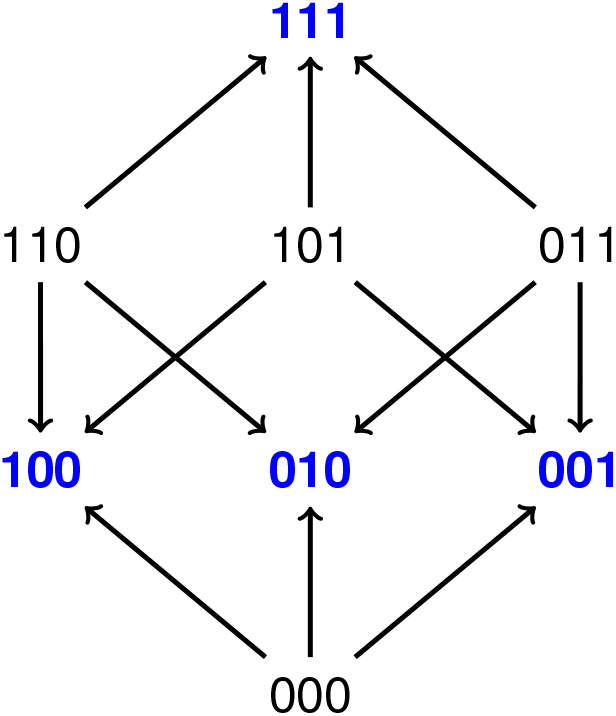
The graph represents fitness landscapes with four peaks. Each arrow points toward the genotype with higher fitness. A biallelic 3-locus system can have at most three peaks in the absence of higher order epistasis.

**Figure 2.**
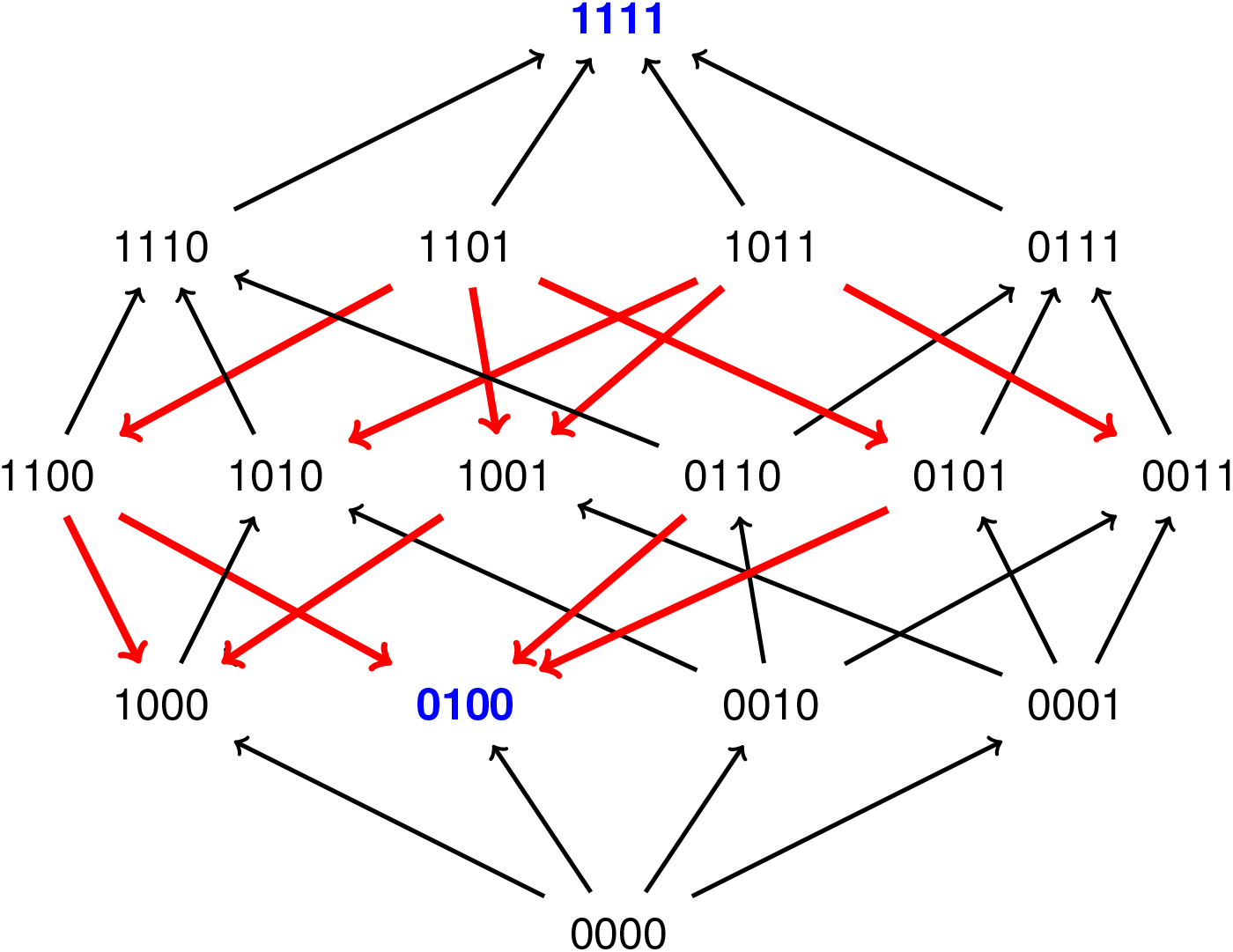
The 16 genotypes represent all combinations of four mutations that individually increase resistance to the malaria drug pyrimethamine in *Plasmodium vivax* (Ogbunugafor and Hartl, 2016). The genotype 1111 is the global peak, `0100` a local peak, and the wild-type 0000 has the lowest fitness in the drug environment.

**Figure 3.**
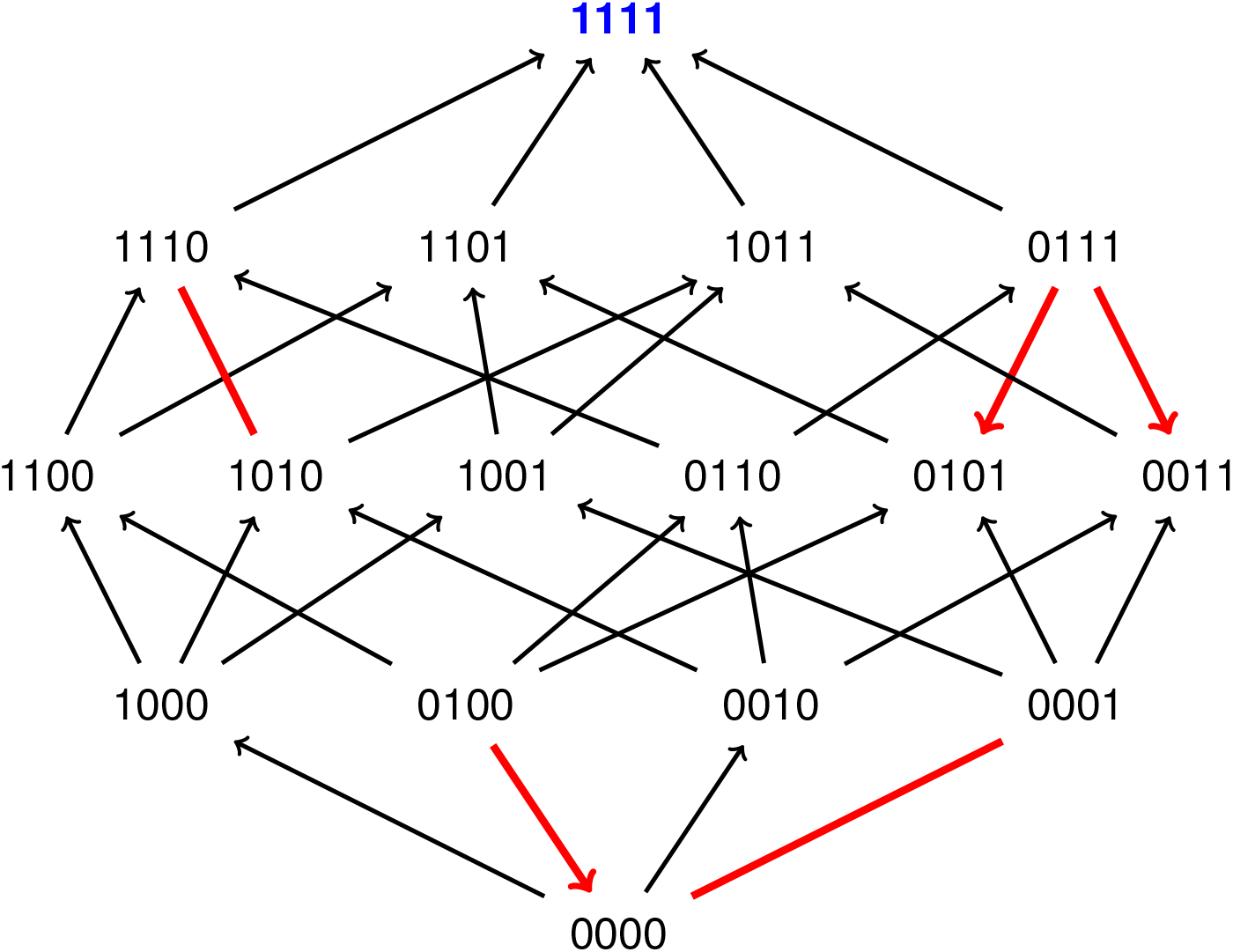
The fitness graph for the wild-type TEM-1, denoted `0000`, and all combinations of the four mutations G238S (1000), M182T (0100), E104K (0010), and A42G (0001). Each arrow points toward the genotype with higher fitness and line segments (marked red) indicate equal fitness. The landscape is single peaked. However, some four-step trajectories to the peak are inaccessible because of the red arrows and line segments.

On the empirical side, experimental studies show that higher order epistasis can shape accessible trajectories (Sailer and Harms, 2017). Constraints on the order in which mutations accumulate have been observed in both experimental and clinical settings. For instance, the pair of mutations G238S and M182T on a TEM-1 background confers cefotaxime resistance, with the evolutionary constraint that G238S has to be in place before M182T is selected for (Singh and Dominy, 2012; Weinreich et al., 2006). Similar order constraints have been reported in viral and parasitic drug resistance, including HIV and malaria, and analyzed using mutational tree models and conjunctive Bayesian networks (Beerenwinkel et al., 2007b; Desper et al., 2000). Some of these constraints require higher-order epistasis.

A signed interaction is defined as a gene interaction that can be deduced from the rank order of genotypes alone. The apparent disagreement between empirical observations of signed interactions and the claimed simplicity of some approaches that use global epistasis cannot be dismissed as an artifact. Rank orders are invariant under sigmoid transformations, and similarly, any type of global epistasis approach that preserves rank orders is expected to detect a considerable fraction of higher order interactions that are signed.

The antibiotic resistance study in Figure 3 has been central to this discussion (Weinreich et al., 2006). It has been cited both in support of signed gene interactions that limit peak accessibility and of a lack of substantial higher order effects. Its chemistry has also been characterized independently (Singh and Dominy, 2012), providing a complementary perspective grounded in protein folding mechanisms.

Here we perform a comprehensive study of signed interactions in the antibiotic resistance and malaria studies (Figures 2 and 3). In addition to sign epistasis in the conventional sense (Weinreich et al., 2005), and signed higher order interactions (Crona et al., 2017), we introduce signed bipyramids as measures comparing extreme and intermediate genotypes along a fitness axis. The bipyramid measures are particularly transparent, as they do not average over different backgrounds, but rather capture local features of the landscape in a direct way.

We then compare the results of the signed analysis with the statistical findings of Park et al. (2024). The signed analysis points out higher order interactions in a specific region of the landscape. Accordingly, we perform a statistical analysis, and confirm substantial higher order interactions in the local setting. A closer examination reveals pronounced scale heterogeneity, which is unsurprising in the context of drug resistance. Signed measures are robust to scale heterogeneity, i.e., they can detect interactions that are locally significant even if they contribute only marginally to a global average. This property helps clarify the dynamics for these landscapes, in particular the role of higher order interactions.

## 2. Results

### 2.1. Signed interactions for antimicrobial drug resistance

A fitness graph of a biallelic *L*-locus system is a directed cube graph where the vertices represent genotypes and each arrow is directed toward the genotype of higher fitness (Crona et al., 2013; De Visser et al., 2009). By convention, the wild-type is denoted by zeros and is placed as the lowest point in the figure.

Sign epistasis means that the effect of a mutation, whether positive or negative, depends on genetic background. Analogously, signed interactions can be inferred from the rank order of genotypes alone. The total third-order epistasis is measured by the form

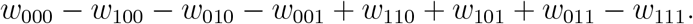

By definition, the system has a third-order interaction if the form is not zero, and a signed third-order interaction if the rank order of genotypes implies that the form is not zero. Note that genotypes with an even number of 1’s have positive coefficients, and the other coefficients are negative.

The fitness graph in Figure 1 shows that there is higher order epistasis in the system, since the odd genotypes `111, 100, 010, 001` are peaks, and the other genotypes are sources in the graph. In fact, already the number of peaks (marked blue) implies that the system has higher order epistasis. A characterization of fitness graphs that require higher order epistasis is given in Crona et al. (2020); Lienkaemper (2018), see also Methods.

The number of peaks and signed third-order interactions are relevant for assessing how consequential the gene interactions are in a system. However, such statistics can be rather coarse, especially for small systems where the upper bounds for these characteristics are small. Rank orders carry information of other types. In cases where there is a clear axis in the *L*-cube along which fitness increases, it can be instructive to consider bipyramid interactions aligned with this axis. For instance, if fitness is expected to increase for mutations 0 →1 in a biallelic 3-locus system, then a bipyramid form compares the triple mutant and the wild type against the three corresponding single mutants, i.e. the form is

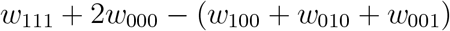

Note that the form is zero for additive fitness.

Figure 2 illustrates a study of malaria drug resistance. The study concerns all combinations of four mutations that confer resistance to pyrimethamine in *Plasmodium vivax* (Ogbunugafor and Hartl, 2016).

Figure 3 illustrates a study on antibiotic resistance, specifically TEM-1 *β*-lactamase. The line segments in the graph indicate equal fitness. The original study (Weinreich et al., 2006) includes all combinations of M182T, G238S, E104K, and A42G on TEM-1 background, as shown in Figure 3, and also a fifth mutation, g4205, not represented in Figure 3. The minimum inhibitory concentration (MIC) values were measured for all genotypes.

The genotype `1111` has the highest fitness in both Figures 2 and 3, whereas the genotype `0000` has the lowest fitness in Figure 3, and second-lowest fitness in Figure 2. Consequently, it is justified to consider bipyramids along the axis from `0000` to `1111`.

In summary, the malaria study has two peaks, two signed third-order interactions and 10 bipyramid interactions out of 16 possible (Table 4). The TEM-1 study is single peaked, has one signed third-order interaction, and one bipyramid interaction. Explicit descriptions of the signed interactions are provided in Methods. Intuitively, the fitness graphs agree well with this summary, in that the malaria landscape appears more “hostile”.

A potential concern about statistics based on signed approaches is significance. However, identifying a signed third-order interaction requires consistent results across four pairwise comparisons, which makes the conclusions relatively robust to moderate levels of error. If measurements are available, significance can be analyzed as conventional. Signed approaches can also be applied in the absence of direct measurements, for instance in strain competition data, where the underlying distribution cannot be directly inferred. In such cases, the error rate in pairwise comparisons is useful.

We performed simulations to relate error rates in pairwise comparisons to incorrectly inferred higher order interactions. Assuming normal distributions for both fitness values and measurement errors, we found that a 10 percent error rate in pairwise comparisons corresponds to a false positive rate of approximately one percent for third-order interactions (see Methods for the detailed assumptions).

### 2.2. A case study: TEM-1 *β*-lactamase

Motivated by the different interpretations of gene interactions for the TEM-1 study, we performed a more detailed analysis. Recall that the global 5-locus system consists of all combinations of M182T, G238S, E104K, A42G, and g4205, whereas Figure 3 shows the 4-locus system consisting of all combination of these mutations with the exception of g4205. A striking property of the global 5-locus system is that mutations vary substantially in how consistent their effects are. As noted by the authors of the original study, the fractions of positive effects over all 16 backgrounds are:

M182T `8/16`, G238S `16/16`, E104K `15/16`, A42G `12/16`.

The described pattern is, to some extent, reflected in Figure 3. Table 2 provides MIC values (left column) and ranks (middle column) for the genotype in Figure 3. Three out of five red arrows and line segments are related to the effect of the substitution M182T, represented by the second locus (from the left). The inconsistent effect of M182T should not be mistaken for weak effects. In fact, each one of the four TEM-mutations increases the MIC-value by at least a factor 10 on some genetic background (Table 3).

Another striking pattern in Figure 3 is that 4 out of 5 red arrows and line segments concern the subsystem consisting of all combinations of M182T, E104K, and A42G on the wild-type background, i.e., the 8 genotypes obtained by varying these three loci while keeping all other loci fixed at the wild-type alleles (corresponding to genotypes whose label begins with a zero). This is also the only subsystem in the graph with a signed third-order interaction.

The MIC values for the genotypes in this subsystem (Table 2, left column) indicate that this signed third-order interaction is substantial, a conclusion that also holds when switching to a log scale.

We performed a statistical analysis for comparing the third-order interactions observed here with those reported in Park et al. (2024). Following the approach in Park et al. (2024), a sigmoid link function was applied (see Methods). We then computed interaction terms up to third-order for this subsystem using the code provided by the authors. In this isolated setting, the third-order effect exceeds most lower-order effects (Table 1, left column).

**Table 1.** Effects on MIC values for all combinations of M182T, E104K and A42G on TEM-1 background. The intercept is denoted ∅ . The left column describes the intercept, main effects, pairwise effects and the third-order effect for the 3-locus system. The non-specific parameters for the system are *L* = −1.1502 and *U*− *L* = 3.2683. The third-order effect (highlighted) exceeds the pairwise and main effects, with a single exception. For easy comparison, the effects in the right column concern the entire 5-locus system that combines five mutations, including M182T, E104K and A42G (the values agree with Table 6).

| Loci | Effects in 3-locus system | Effect in 5-locus system (from Table 2) |
| --- | --- | --- |
| ∅ | -0.77612 | -2.0653 |
| 1 | 0.2386 | 0.13452 |
| 2 | 0.30475 | 0.32731 |
| 3 | 1.0113 | 0.20746 |
| 12 | -0.56267 | 0.067337 |
| 13 | 0.22887 | 0.011705 |
| 23 | 0.16273 | 0.14171 |
| 123 | 0.6109 | -0.024313 |

**Table 2.** MIC values and rank orders for all combinations of the mutations G238S (1000), M182T (0100), E104K (0010), A42G (0001). The rightmost column shows ranks based on estimates from at most pairwise effects. The ranks are boldfaced for the wild-type and its neighbors.

| Sequence | MIC | Rank from measurements | Rank from estimates |
| --- | --- | --- | --- |
| 0000 | 0.088 | **14** | **8** |
| 1000 | 1.4 | **8** | **12** |
| 0100 | 0.063 | **16** | **13** |
| 0010 | 0.13 | **13** | **14** |
| 0001 | 0.088 | **14** | **9** |
| 1100 | 32 | 6 | 11 |
| 1010 | 360 | 3 | 4 |
| 1001 | 23 | 7 | 6 |
| 0110 | 0.18 | 12 | 16 |
| 0101 | 1.4 | 8 | 15 |
| 0011 | 1.4 | 8 | 7 |
| 1110 | 360 | 3 | 3 |
| 1101 | 360 | 3 | 5 |
| 1011 | 2100 | 2 | 2 |
| 0111 | 0.8 | 11 | 10 |
| 1111 | 2900 | 1 | 1 |

**Table 3.**
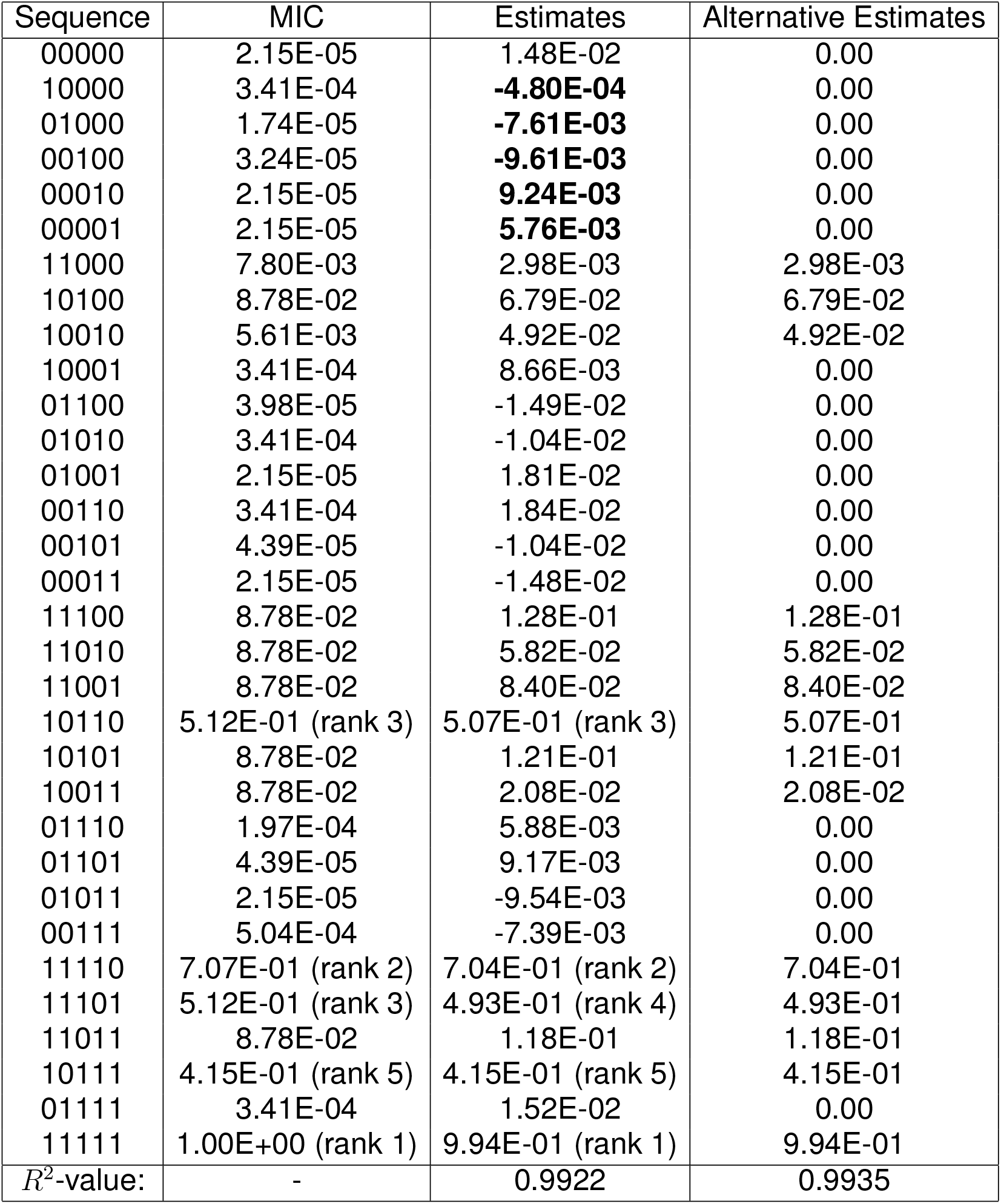
The first column provides MIC values (normalized, so that the highest value equals 1 rather than the original value 4100). The second column gives estimates based on at most pairwise interactions. The third column is an alternative estimate obtained by copying the second column, except that entries are set to 0 whenever the normalized MIC value is ≤3.41 ·10^*−*4^. The boldface estimates concern neighbors of the wild-type, all of them lower than the wild-type estimate (which would prevent evolution). The estimates for high MIC values agree well with measurements, in particular the ranks are nearly perfect for the five genotypes with the highest MIC values (with ranks provided in paranthesis). The bottom of the table shows *R*^2^-values for the estimates and alternative estimates.

**Table 4.**
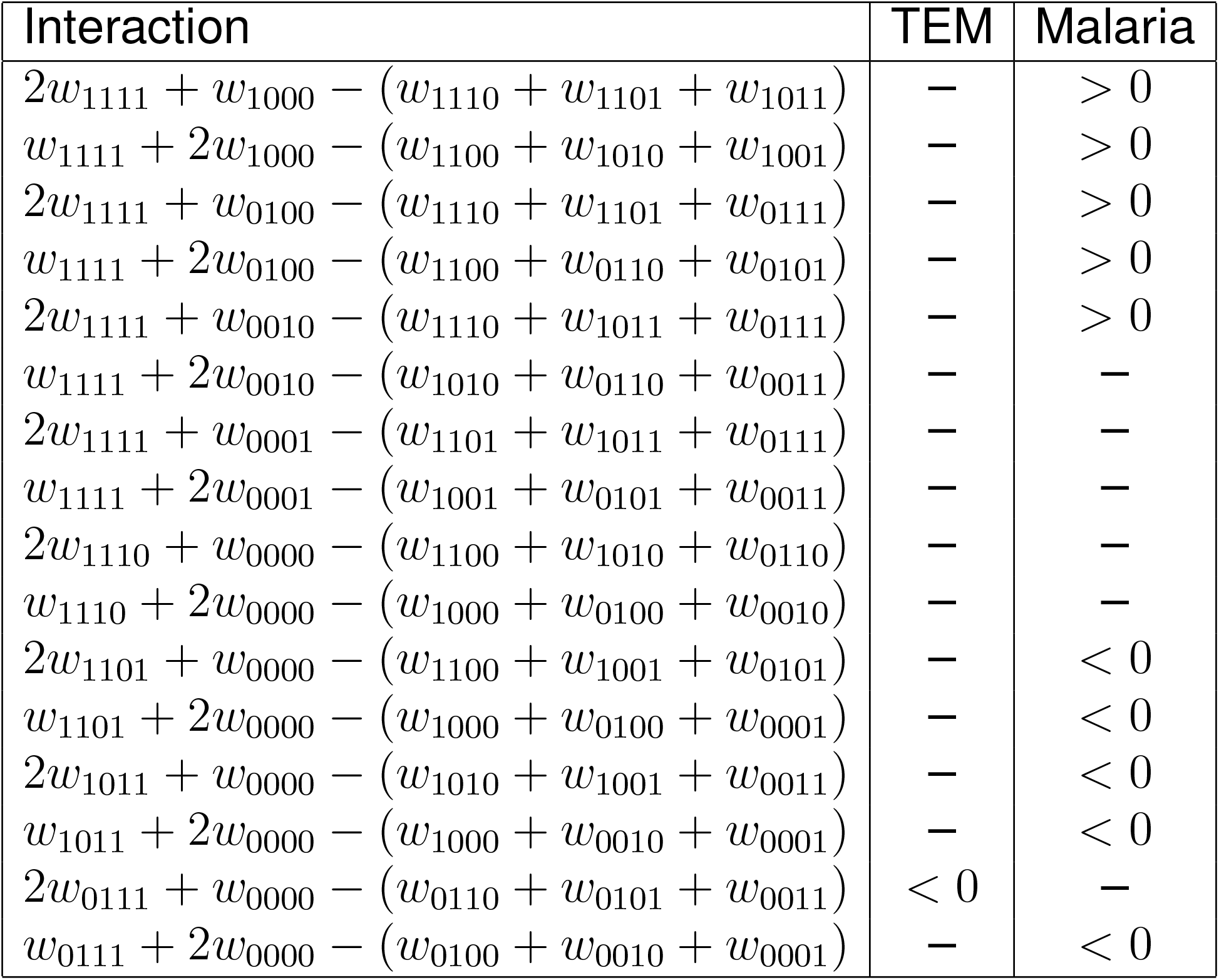
Bipyramid circuit interactions.

Table 1 compares the effect for this subsystem in isolation against the corresponding result in the global 5-locus system Park et al. (2024). In the global system, the third-order effect is much smaller than lower-order effects. The third-order effects in both settings are highlighted in the table for direct comparison.

The observations in Table 1 can be compared by the MIC values for the global 5-locus system. The genotypes that combines M182T, E104K, and A42G on the wild-type background have MIC value ≤ 1.4, whereas the same mutations in the the presence of g4205 have MIC values up to 2100 (Table 3, left column, provides normalized MIC-values). As a result, the contribution of this subsystem on the wild-type background is effectively suppressed when averaged over all genetic backgrounds.

Similar to the results in Table 1, the third-order effects are smaller than the lower-order effects (Table 6) in the 5-locus system. Because of the noted scale heterogeneity, it can be instructive to study rank orders according to estimates based on at most pairwise interactions following Park et al. (2024). These rank orders are compared to rank orders according to measurements in Table 2. The table shows disagreements for the wild-type neighbors. If the estimate would be correct, then evolution from TEM-1 would not be possible under standard assumptions, since TEM-1 would be a fitness peak.

For an additional perspective, one can consider the underlying chemistry. An independent study (Singh and Dominy, 2012) shows that G238S and E104K (about 10 A away from the active site) increase Cefotaxime hydrolysis, whereas M182T and A42G (about 20 A away from the active site) do not, in agreement with the MIC measurements. It has been established that G238S and E104K directly restructure the active site while M182T acts as a stabilizer (Singh and Dominy, 2012; Wang et al., 2002) (stabilizers can reduce misfolding and their benefits are context dependent). The independent observations seem compatible with the rank order observations according to measurements, i.e., the 16/16 positive effects for G238S, 15/16 for E104K, and 8/16 for M182T. In contrast, none of the mutations A42G, E104K, M182T, or G238S increase fitness according to the estimate based on at most pairwise interactions (Table 2).

Further comparisons between estimates based on at most pairwise effects and measurements are provided in Table 3. From the table, the agreement appears nearly perfect for the five genotypes with highest measured MIC values, whereas there seems to be a disagreement for genotypes with low MIC values.

For testing the hypothesis that the disagreement is substantial for low MIC values, we defined an alternative estimate by assigning the value zero to approximately half of the genotypes, specifically the 18 genotypes with lowest MIC values (Table 3). All values for the alternative estimates were identical to those of the original estimate. The *R*^2^ value for the alternative estimate was similar, in fact marginally higher, than that of the estimate itself. This result suggests that for these 18 genotypes the estimates provide little information about the measurements beyond indicating that the measured values are small.

## 3. Discussion

The importance of higher order interactions remains disputed. Several unrelated arguments in the older literature have suggested that it is sufficient to consider at most pairwise epistasis (see Crona et al. (2017) for a comment on the history, and also Crona (2016)). Empirical support for the opposite view has been provided in (Weinreich et al., 2018; Weinrech et al., 2013), and recent discussions include (Park et al., 2024b; Dupic et al., 2024; Park et al., 2024).

This question is of general interest because higher order epistasis can strongly shape fitness landscapes and, consequently, evolutionary dynamics. As shown previously, there is an association between higher order epistasis and the number of peaks (Crona et al., 2020); for instance, the fitness graph in Figure 1, with four peaks, is incompatible with its absence. Higher order epistasis can also serve as an indicator of complex chemistry. From a practical perspective, its presence affects the ability to predict unseen genotypes.

A recent approach argues that the protein-sequence-function relationships can be described in a simple way if one allows for global epistasis (Park et al., 2024). Park and coauthors have developed an approach that applies a sigmoid link function to capture global nonlinearities. For a collection of empirical studies, the authors show that at most pairwise interactions, together with a global nonlinearity, explain at least 92 percent of the phenotypic variance in the studied systems.

However, rank order based studies indicate that higher order interactions are present in many fitness landscapes (Crona et al., 2017). Whenever the rank order of genotypes implies that gene interactions are present, these interactions are defined to be signed.

One study of antibiotic resistance has been cited for rich signed interactions that limit the number of evolutionarily accessible trajectories to the peak, and is also included in the collection from (Park et al., 2024). The study concerns all combinations of four clinically found TEM-1 *β*-lactamase mutations E104K, G238S, A42G, and M182T (Weinreich et al., 2006), see (Figure 3). By inspection, the figure reveals a signed third-order interaction according to general theory on partial orders and interactions (Crona et al., 2017; Lienkaemper, 2018).

To resolve this apparent contradiction, we performed a detailed case study and found that scale heterogeneity appears to explain the discrepancy. The signed third-order interaction was substantial in relative terms, i.e., it exceeded most lower order effects in its 3-locus subsystem. This interaction impacts evolutionary trajectories and also reflects the underlying biochemistry, specifically the inconsistent effect of the stabilizing mutation M182T. However, the corresponding third-order effect was very small at the global system level.

To investigate the role of scale further, we systematically compared estimates based on global epistasis and pairwise interaction with the direct measurements. The agreement is very good for high-fitness genotypes. However, the estimated rank order among low-fitness genotypes appears more or less unrelated to the rank order based on measurements. In particular, evolution from the wild-type would not be possible according to the estimate, as the wild-type is ranked higher than all its neighbors. In contrast, the wild-type has second-lowest rank according to measurements. As an additional check, an alternative estimate was constructed that was identical to the original estimate, except that approximately half of the genotypes with the lowest fitness were assigned the value of zero. For the alternative estimate, the *R*^2^ value is similar, in fact marginally higher, than for the estimate itself. This outcome suggests that the original estimate does not capture information about the measurements for these genotypes, beyond indicating that their values are low. These observations support that scale heterogeneity is a major factor.

One can ask if features of the TEM-1 study are present in other systems. In antimicrobial resistance, the wild-type tends to have low fitness in a drug environment, whereas resistant genotypes can attain proxy measures that are higher by factors of thousands. Consequently, approaches that ignore these effects are likely to underestimate the impact of higher order interactions. Regression methods in a broad sense, including Walsh coefficients, with or without global epistasis, may be sensitive to this effect.

Signed measures can be used as a complement to other methods. Their advantages include invariance under many transformations and robustness to scale heterogeneity. An obvious limitation is that they do not capture gene interactions that do not affect rank orders.

The signed interactions in the TEM-1 study are by no means exceptional. For comparison, the study of malaria-drug resistance (Figure 2) is richer in signed third-order interactions, as well as in signed bipyramid interactions (Table 5), and has more peaks. The collection of empirical studies in (Park et al., 2024) includes a second study of antibiotic resistance, specifically on dihydrofolate reductase. Without performing a detailed analysis, we noted that 21 biallelic subsystems out of 96 3-locus subsystems, or about 22 percent, show signed third-order interactions. Several empirical sets with signed higher order interactions are also discussed in Crona et al. (2017).

**Table 5.** MIC values and rank order for all combinations of the mutations G238S (10000), M182T (01000), E104K (00100), A42G (00010), g4205 (00001). For the reader’s convenience, the rightmost column provides ranks within the subsystem consisting of all combinations of the three mutations M182T, E104K and A42G. The corresponding MIC values are bold-faced.

| Sequence | MIC | Rank | Rank in subsystem |
| --- | --- | --- | --- |
| 00000 | **0.088** | 13 | 5 |
| 10000 | 1.4 | 9 | - |
| 01000 | **0.063** | 14 | 6 |
| 00100 | **0.13** | 12 | 4 |
| 00010 | **0.088** | 13 | 5 |
| 00001 | 0.088 | 13 | - |
| 11000 | 32 | 6 | - |
| 10100 | 360 | 5 | - |
| 10010 | 23 | 7 | - |
| 10001 | 1.4 | 9 | - |
| 01100 | **0.18** | 11 | 3 |
| 01010 | **1.4** | 9 | 1 |
| 01001 | 0.088 | 13 | - |
| 00110 | **1.4** | 9 | 1 |
| 00101 | 0.18 | 11 | - |
| 00011 | 0.088 | 13 | - |
| 11100 | 360 | 5 | - |
| 11010 | 360 | 5 | - |
| 11001 | 360 | 5 | - |
| 10110 | 2100 | 3 | - |
| 10101 | 360 | 5 | - |
| 10011 | 360 | 5 | - |
| 01110 | **0.8** | 10 | 2 |
| 01101 | 0.18 | 11 | - |
| 01011 | 0.088 | 13 | - |
| 00111 | 2.0 | 8 | - |
| 11110 | 2900 | 2 | - |
| 11101 | 2100 | 3 | - |
| 11011 | 360 | 5 | - |
| 10111 | 1500 | 4 | - |
| 01111 | 1.4 | 9 | - |
| 11111 | 4100 | 1 | - |

**Table 6.** Effects on MIC values for all combinations G238S (10000), M182T (01000), E104K (00100), A42G (00010) and (00001) on TEM-1 background. The non-specific parameters are *L* = − 0.12861 and *U* −*L* = 1.7701. The intercept is denoted ∅ and the effects for each combination of up to three loci are listed.

| Loci | Effect |
| --- | --- |
| ∅ | -2.0653 |
| 1 | 0.49036 |
| 2 | 0.13452 |
| 3 | 0.32731 |
| 4 | 0.20746 |
| 5 | 0.087472 |
| 12 | 0.13673 |
| 13 | 0.32829 |
| 14 | 0.20586 |
| 15 | 0.084244 |
| 23 | 0.067337 |
| 24 | 0.011705 |
| 25 | 0.078903 |
| 34 | 0.14171 |
| 35 | 0.024871 |
| 45 | -0.038974 |
| 123 | -0.073047 |
| 124 | -0.0012334 |
| 125 | -0.076443 |
| 134 | -0.12697 |
| 135 | -0.02083 |
| 145 | 0.031844 |
| 234 | -0.024313 |
| 235 | -0.11107 |
| 245 | 0.050073 |
| 345 | 0.036059 |

It follows directly that allowing for global epistasis by using any type of link function reduces the contributions of other forms of interactions. It is likewise clear that taking global non-linearities into account, if such effects are present, can lead to a simpler and more natural description of data. However, it does not follow that higher order interactions are absent in biological systems, or that they fail to carry important information.

As an additional complication, a simple sequence-phenotype relationship does not imply a simple sequence-fitness relationship (Srivastava and Payne, 2022). For example, even if mutations contribute additively to a phenotype, selection may strongly favor phenotypic values that exceed a certain threshold. Such nonlinear selection can induce higher-order epistasis in the resulting fitness landscape. Related threshold dynamics are discussed in Dupic et al. (2024).

Pairwise interactions have been reported even in very large fitness landscapes. In some cases, two recurrent mutations in an adapting population almost never co-occur. For antibiotic resistance, the clinically important TEM-1 mutations G238S and R164 rarely co-occur. For SARS-CoV-2, mutations at positions N450 and R346 have never been observed together in Omicron genomes, despite the analysis of millions of sequences worldwide (Focosi et al., 2023).

In contrast, clear empirical evidence for consistent third-order interactions, such as mutation triples that are systematically favored or disfavored beyond pairwise effects in large fitness landscapes, remains limited. Whether consistent higher order interactions can be important in large fitness landscapes remains an open question.

## 4. Methods

### 4.1. Signed third-order interactions

A biallelic 3-locus system consists of eight genotypes

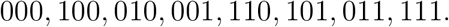

The total third-order epistasis is measured by the form

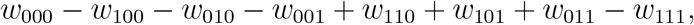

or

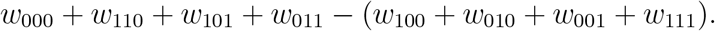

Note that genotypes with an odd number of 1’s are compared with the genotypes with an even number of 1’s, which we denote *o* and *e*, respectively. By definition, the system has a third-order interaction if the form is not zero, and a signed third-order interaction if the rank order of genotypes implies that the form is not zero.

One can phrase an exact criterion for signed third-order interactions that applies to fitness graphs and any other partial orders:

A partial order ≺ of the genotypes with respect to fitness implies 3-way epistasis if and only if there exist a partition {*o*_*j*_, *e*_*j*_} of all genotypes such that

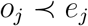

for all *j*, or vice versa. The analogous criterion holds for epistasis of any order (Crona et al., 2020; Lienkaemper, 2018).

Figure 1 has a signed third-order interaction, since the genotypes `111, 100, 010` and `001` are sinks (peaks), whereas the remaining genotypes are sources.

Exactly two subsystems for the malaria study (Figure 2) have signed third-order epistasis. One of them is {`1110, 1100, 1010, 0110, 1000, 0100, 0010`} . Notice that the fourth coordinate is zero for all genotypes, so that the coordinate is simply a placeholder. The signed interaction follows from the inequalities

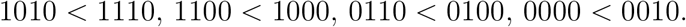

The second subsystem is {`1111, 1110, 1101, 1011, 1100, 1010, 1001, 1000`} . The signed interaction follows from the inequalities

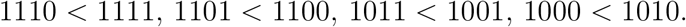

By inspection, no other subsystems have signed third-order interactions.

For the TEM-1 study (Figure 3) the subsystem consisting of the mutations M182T, E104K, and A42G on the wild-type background has a signed higher order interaction. With notation as in Figure 3, the system is

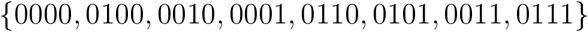

The first coordinate is zero for all eight genotypes in the inequality, indicating the absence of G238S in the subsystem. The signed interaction

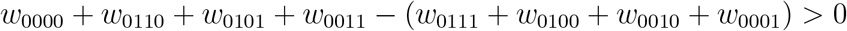

follows from the inequalities.

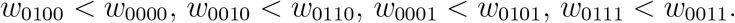

By inspection, no other subsystems in the TEM-1 study have a signed third-order interaction.

### 4.2. Signed bipyramid interactions

As a further comparison between the malaria and TEM-1 studies Table 4 shows bipyramid interactions. Specifically, bipyramids such that the exreme genotypes have distance 3 (differ by three mutations), are considered. It can be instructive to consider an example from the malaria study. Consider the bipyramid with top genotype `1011` and bottom genotype `0000`, and the intermediate double mutants. The bottom genotype `0000` has the lowest rank in the system (16), and the intermediate double mutants have higher ranks 3, 10 and 11. Nevertheless, the rank of `1011` is 14, among the lowest in the system. For the bipyramid discussed, the rank order implies a signed interaction

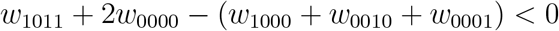

All bipyramid interactions are zero if fitness is additive. The malaria study has 10 bipyramid interactions, and the TEM-1 study has one such interaction (Table 4).

The 10 bipyramid interactions show a consistent pattern. The genotype 1111 has high fitness, relative to expectations based on double and triple mutants, whereas some triple mutants have low fitness, relative to expectations based on single and double mutants.

The rank orders for the malaria study are provided in Table 7, and for the TEM-1 study in Table 5, third column.

**Table 7.** Rank order of Malaria genotypes for all combinations of four mutations.

| Sequence | Rank |
| --- | --- |
| 0000 | 16 |
| 1000 | 5 |
| 0100 | 4 |
| 0010 | 12 |
| 0001 | 15 |
| 1100 | 6 |
| 1010 | 3 |
| 1001 | 11 |
| 0110 | 8 |
| 0101 | 9 |
| 0011 | 10 |
| 1110 | 2 |
| 1101 | 13 |
| 1011 | 14 |
| 0111 | 7 |
| 1111 | 1 |

Bipyramids are circuits, or minimal dependence relations, as defined in matroid theory. Another class of signed circuits is used in (Crona, 2020). Note that bipyramid interactions do not imply higher order epistasis, and the reverse implication does also not hold.

For more context, the complete set of circuits signs for an *L*-cube determines a triangulation of the cube. Such a triangulation is referred to as the shape of the fitness landscape (Beerenwinkel et al., 2007a). For mathematical background, see De Loera et al. (2010). For recent contributions to the theory of signed interactions in evolutionary biology, including statistical approaches, see Ribeca et al. (2026); Carlson et al. (2025); Pahujani and Krug (2025); Krug and Oros (2024); Crona et al. (2023); Riehl et al. (2022); Saona et al. (2022); Das et al. (2020).

### 4.3. Rank order methods and significance

We now consider the effect of measurement noise on detecting signed interactions. Assume that a fitness proxy is normally distributed with standard deviation *σ*, and that the measurement errors are normally distributed with standard deviation 0.23*σ*.

According to simulations, the success rate for pairwise comparison is 90 percent under these assumptions.

To identify a third-order interaction, such as

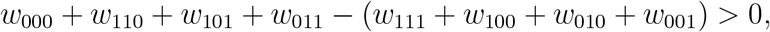

one needs to compare four measured values *a*_1_, *a*_2_, *a*_3_, *a*_4_ for the even genotypes with four measured values *b*_1_, *b*_2_, *b*_3_, *b*_4_ for the odd genotypes. For simplicity, we condition on the event that the observed values are correctly ranked *a*_1_ *> a*_2_ *> a*_3_ *> a*_4_ and *b*_1_ *> b*_2_ *> b*_3_ *> b*_4_.

One infers the third-order interaction above exactly if *a*_*i*_ *> b*_*i*_ for all *i*. The conclusion is considered correct if

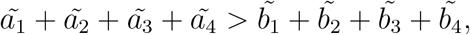

where 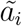 and 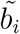 denote the true (noise-free) values. Simulations based on 10,000 experiments indicate that the false positive rate is approximately one percent.

### 4.4. Global epistasis

By using a sigmoid link function one can estimate global non-linearities in data (Park et al., 2024). In brief, a lower bound *L*, an upper bound *U*, and effects of an arbitrary number of orders are inferred from data. For instance, an estimate of the fitness *y*(*g*) based on at most pairwise effects is

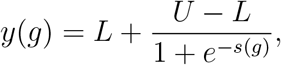

where *s*(*g*) includes main effects and pairwise effects, but no higher order terms.

## Notes

### Competing Interest Statement

The authors have declared no competing interest.

### Summary of Updates

More references have been included, along with some background.

## References

Beerenwinkel, N., Eriksson, N., and Sturmfels, B. (2007). Conjunctive bayesian networks. Bernoulli 13 (4)

Beerenwinkel, N., Pachter, L., and Sturmfels, B. (2007). Epistasis and shapes of fitness landscapes. Statistica Sinica, 1317–1342.

Carlson, M. O., Andrews, B. L., and Simons, Y. B. (2025). Distinguishing direct interactions from global epistasis using rank statistics. Proceedings of the National Academy of Sciences, 122(39), e2509444122.

Crona, K. (2016). Epistasis and entropy. PLoS Genetics, 12(12), e1006322.

Crona, K. (2020). Rank orders and signed interactions in evolutionary biology. Elife, 9, e51004.

Crona K., Gavryushkin, A., Greene, D., and Beerenwinkel, N. (2017). Inferring genetic interactions from comparative fitness data. Elife, 6, e28629.

Crona, K., Greene, D. and Barlow, M. (2013). The peaks and geometry of fitness landscapes. Journal of theoretical biology, 317, 1–10.

Crona, K., Krug, J., and Srivastava, M. (2023). Geometry of fitness landscapes: peaks, shapes and universal positive epistasis. Journal of Mathematical Biology, 86(4), 62.

Crona, K., Luo, M., and Greene, D. (2020). An uncertainty law for microbial evolution. Journal of Theoretical Biology, 489, 110155.

Das, S. G., Direito, S. O., Waclaw, B., Allen, R. J., and Krug, J. (2020). Predictable properties of fitness landscapes induced by adaptational tradeoffs. Elife, 9, e55155.

De Loera, J., Rambau, J., and Santos, F. (2010). Triangulations: structures for algorithms and applications (Vol. 25). Springer Science and Business Media.

De Visser, J. A. G., Park, S. C., and Krug, J. (2009). Exploring the effect of sex on empirical fitness landscapes. The American Naturalist, 174(S1), S15–S30.

Desper, R., Jiang, F., Kallioniemi, O. P., Moch, H., Papadimitriou, C. H., and Schäffer, A. A. (2000). Distance-based reconstruction of tree models for oncogenesis. Journal of Computational Biology, 7(6), 789–803.

Dupic, T., Phillips, A. M., and Desai, M. M. (2024). Protein sequence landscapes are not so simple: on reference-free versus reference-based inference. bioRxiv.

Focosi, D., Quiroga, R., McConnell, S., Johnson, M. C., and Casadevall, A. (2023). Convergent evolution in SARS-CoV-2 spike creates a variant soup from which new COVID-19 waves emerge. International journal of molecular sciences, 24(3), 2264.

Kryazhimskiy, S., Rice, D. P., Jerison, E. R., and Desai, M. M. (2014). Global epistasis makes adaptation predictable despite sequence-level stochasticity. Science, 344(6191), 1519–1522.

Krug, J., and Oros, D. (2024). Evolutionary accessibility of random and structured fitness landscapes. Journal of Statistical Mechanics: Theory and Experiment, 2024(3), 034003.

Lienkaemper, C., Lamberti, L., Drain, J., Beerenwinkel, N., and Gavryushkin, A. (2018). The geometry of partial fitness orders and an efficient method for detecting genetic interactions. Journal of Mathematical Biology, 77(4), 951–970

Mira, P. M., Crona, K., Greene, D., Meza, J. C., Sturmfels, B., and Barlow, M. (2015). Rational design of antibiotic treatment plans: a treatment strategy for managing evolution and reversing resistance. PloS one, 10(5), e0122283.

Nichol, D., Rutter, J., Bryant, C., Hujer, A.M., Lek, S., Adams, M.D., Jeavons, P., Anderson, A.R., Bonomo, R.A. and Scott, J.G., (2019). Antibiotic collateral sensitivity is contingent on the repeatability of evolution. Nature communications, 10(1), 334.

Otwinowski, J., McCandlish, D. M., and Plotkin, J. B. (2018). Inferring the shape of global epistasis. Proceedings of the National Academy of Sciences, 115(32), E7550–E7558.

Ogbunugafor, C. B. and Hartl, D. (2016). A pivot mutation impedes reverse evolution across an adaptive landscape for drug resistance in Plasmodium vivax. Malaria journal 15(1), 1–10.

Pahujani, S., and Krug, J. (2025). Complexity and accessibility of random landscapes. SciPost Physics Lecture Notes 108.

Park, Y., Metzger, B. P., and Thornton, J. W. (2024). On the Analysis of Protein Genetic Architecture: Response to “Protein sequence landscapes are not so simple”. bioRxiv.

Park, Y., Metzger, B. P., and Thornton, J. W. (2024). The simplicity of protein sequence-function relationships. Nature Communications, 15(1), 7953.

Ribeca, P., Castro, A., Lage-Castellanos, A., Sergeeva, A., Matuszewski, S., Paccosi, R.G., Belik, V., Ghafari, M., Krug, J., Achaz, G. and Ferretti, L. (2026). Simple sign epistasis and evolutionary detours in fitness landscapes. arXiv preprint arXiv:2604.22611.

Srivastava, M., and Payne, J. L. (2022). On the incongruence of genotype-phenotype and fitness landscapes. PLoS computational biology 18(9), e1010524.

Riehl, M., Phillips, R., Pudwell, L., and Chenette, N. (2022). Occurrences of reciprocal sign epistasis in single-and multi-peaked theoretical fitness landscapes. Journal of Physics A: Mathematical and Theoretical, 55(43), 434002.

Sailer, Z. R. and Harms, M. J. (2017). High-order epistasis shapes evolutionary trajectories. PLoS computational biology, 13(5), e1005541.

Saona, R., Kondrashov, F. A., and Khudiakova, K. A. (2022). Relation between the number of peaks and the number of reciprocal sign epistatic interactions. Bulletin of Mathematical Biology, 84(8), 74.

Singh, M. K., and Dominy, B. N. (2012). The evolution of cefotaximase activity in the TEM β-lactamase. Journal of molecular biology, 415(1), 205–220.

Starr, T. N., and Thornton, J. W. (2016). Epistasis in protein evolution. Protein science, 25(7), 1204–1218.

Wang, X., Minasov, G., and Shoichet, B. K. (2002). Evolution of an antibiotic resistance enzyme constrained by stability and activity trade-offs. Journal of molecular biology, 320(1), 85–95.

Weinreich, D. M., Lan, Y., Jaffe, J., and Heckendorn, R. B. (2018). The influence of higher-order epistasis on biological fitness landscape topography. Journal of statistical physics, 172(1), 208–225.

Weinreich, D. M., Lan, Y., Wylie, C. S., and Heckendorn, R. B. (2013). Should evolutionary geneticists worry about higher-order epistasis? Current opinion in genetics & development, 23(6), 700–707.

Weinreich, D. M., Delaney, N. F., DePristo, M. A., and Hartl, D. L. (2006). Darwinian evolution can follow only very few mutational paths to fitter proteins. Science, 312(5770), 111–114.

Weinreich, D. M., Watson, R. A., and Chao, L. (2005). Perspective: sign epistasis and genetic constraint on evolutionary trajectories. Evolution, 59(6), 1165–1174.

Zhou, J., Wong, M. S., Chen, W. C., Krainer, A. R., Kinney, J. B., and McCandlish, D. M. (2022). Higher-order epistasis and phenotypic prediction. Proceedings of the National Academy of Sciences, 119(39), e2204233119.

